# Are they funny? Associations between instructors’ humor and student emotions in undergraduate lab courses

**DOI:** 10.1101/2025.06.11.659171

**Authors:** Zarae A. Allen, Christopher James Zajic, Christina M. Leckfor, Erin L. Dolan, Trevor T. Tuma

## Abstract

Instructor use of humor can positively affect students’ educational experiences by increasing students’ comfort in the classroom and making the instructor seem more approachable. Humor can also elicit emotional responses, which in turn may influence students’ engagement in the course and their relationship with the instructor. However, students may interpret instructor humor differently, resulting in varied effects. The present study examined how instructors’ use of humor relates to students’ emotions about their lab course. In addition, we examined whether researcher-identified humor aligned with student reports to support valid inferences about instructor humor. To accomplish this, trained researchers analyzed classroom audio recordings of instructor talk in 48 lab courses to identify instances of instructor humor. We also surveyed their undergraduate students (*n* = 462) about their instructor’s humor and pleasant and unpleasant emotions about their lab course. Our results revealed that trained researchers’ coding of instructor humor was poorly predictive of students’ emotions about their laboratory courses. In contrast, students who perceived their instructor to be humorous reported greater pleasant emotions and fewer unpleasant emotions. Our results suggest that instructor humor from students’ perspective may be influential in how they experience instruction. In addition, student perceptions of instructor humor may be a more useful indicator than researcher observations for studying instructor humor.

## INTRODUCTION

The quality of student-instructor relationships is an important factor in students’ academic engagement, achievement, and motivation ^1–4^. Instructors may attempt to build these relationships through their use of language beyond course content. Instructor talk, defined as “language used by an instructor that is not directly related to the concepts under study but instead focuses on creating the learning environment,” includes efforts to build rapport, explain pedagogical choices, share personal experiences, and unmask science ^5,6^. Instructor talk may shape students’ perceptions of instructor immediacy, or the sense of closeness between instructors and students ^7^. Instructor immediacy behaviors, such as smiling, making eye contact, using students’ names, and incorporating humor, have been associated with students’ motivational beliefs, classroom participation, learning, and academic engagement ^8–14^.

One way instructors may attempt to enhance their immediacy and build stronger relationships with students is the use of humor ^15,16^. Humor refers to a psychological response that can be expressed in one of three ways: laughing (e.g., behavioral), perceiving something as funny (e.g., cognitive), or experiencing amusement (e.g., emotional) ^17,18^. Humor often involves both verbal and non-verbal communication that elicits laughing, chuckling, or joy, such as using funny examples, word play, exaggerated descriptions, and funny anecdotes ^19,20^. Instructors may use humor while teaching to foster a sense of community in the classroom, create a more engaging learning environment, and present themselves as more approachable and relatable to students ^21–23^. While instructor humor can be perceived positively, its impact ultimately depends on how it is received by students. Perceptions of humor are subjective, and some forms of instructor humor may be viewed as unfunny, inappropriate, or alienating, potentially undermining the instructor-student relationship ^5,24,25^. Given humor’s potential to enhance instructor immediacy and support relationship-building, further research is needed to understand how instructor humor shapes the classroom environment and students’ learning experiences.

Undergraduates can experience a variety of emotions, meaning their experiential, physiological and behavioral responses to personally meaningful stimuli ^26^, during their educational experiences ^27–29^. Students’ emotions are consistently associated with their academic achievement, motivation, and study strategies, which highlights the importance of identifying the factors that influence their emotions in the classroom ^30–33^. Humor has the potential to yield a range of emotional responses depending on various contextual factors, such as the content of the humor, social dynamics within the classroom, and individual characteristics that influence students’ educational experiences ^34,35^. Instructor humor can promote an effective student-teacher relationship, foster a positive classroom climate, and influence student emotions ^36,37^. For example, humor that is perceived positively can increase student enjoyment and decrease their unpleasant emotions such as anger, anxiety, and boredom ^22,38–40^. In contrast, humor that is perceived negatively is associated with reduced enjoyment and increased boredom, anxiety, and anger ^19,21,41^.

Lab courses may present distinct opportunities for instructors to be humorous because the class sessions are longer, typically enroll fewer students, and are less predictable than in lecture courses. These elements may present more opportunities for what is known as a “violation” (e.g., something unexpected, awkward, or potentially problematic). Benign violation theory ^42–44^ suggests that events are likely to be perceived as humorous when a situation involves a violation that is perceived as benign (e.g., non-threatening or harmless). For instance, students may make technical errors when conducting lab tasks (e.g., pipetting incorrectly, forgetting to include a control sample in an experiment) and instructors might use lighthearted language or make a joke to help students recognize and address the issue rather than dwelling on it. Instructors may also experience potentially problematic events, such as making a mistake when conducting calculations in front of the class or malfunctions of equipment. By framing these events humorously, the instructor can make them feel less threatening for students and for themselves, promoting more pleasant emotions about the course and greater engagement in the course. However, instructor humor can also backfire when it is perceived by students as not taking the situation, their work, or them seriously ^45,46^. With this in mind, we sought to address the following research question: How does instructor humor relate to student emotions about their lab courses?

To address our research question, we collected audio recordings of instructors as they taught lab courses and surveyed students about their emotions toward their lab course at the conclusion of the term. We used the Circumplex Model of Affect to conceptualize and measure students’ emotions ^47^. This model posits that emotions can be described along two broad dimensions: valence and activation. Valence ranges from pleasant (e.g., happy) to unpleasant (e.g., sad), while activation ranges from high (e.g., alert) to low (e.g., sleepy). Different emotions can be understood as a combination of these two dimensions, or as varying degrees of valence and activation. For example, students may experience emotional states that are high in activation and unpleasantness (e.g., *nervous* to use an expensive piece of equipment), high in activation and pleasantness (e.g., *excited* to perform an experiment), low in activation and unpleasantness (e.g., *sad* they are in lab), or low in activation and pleasantness (e.g., *calm* after submitting an assignment). Thus, we surveyed students to detect any relationships between their instructor’s use of humor and the valence and activation level of their emotions about their lab course. We hypothesized that instructor humor would relate positively to pleasant emotions and negatively to unpleasant emotions.

To address our research question, we also needed to measure instructor humor. Yet, humor is challenging to measure because it is interpreted subjectively and can have varying effects depending on the context and how it is perceived ^24,25^. Existing classroom instruments useful for observing teaching practices are not designed to measure instructor humor ^48–51^. Thus, we sought to measure humor in two ways: systematically reviewing transcripts of class recordings (i.e., researcher observation) and surveying students about the extent to which their instructor was humorous (i.e., student report). Our two-fold approach enabled us to address a second research question: How do researcher observations of instructor humor relate to student perceptions of instructor humor? We hypothesized that researcher observations of instructor humor would correlate positively with student reports of instructor humor.

## METHODS

### Ethical guidelines

All participants were treated in accordance with APA ethical guidelines. This research was part of a larger research study, which was reviewed and determined to be exempt by the University of Georgia’s Institutional Review Board (PROJECT00003103).

### Participants

We collected audio recordings of 48 instructors as they taught introductory life science lab courses, and survey data on students’ emotions about the lab course and the extent to which their instructor was humorous at the conclusion of the term. The data presented were collected as part of a larger study examining instructor talk in undergraduate lab courses ^52^. The present study utilizes a subset of data from the broader dataset and overlaps only in instructor and student demographics.

#### Instructors

We used a purposeful sampling strategy to recruit a national sample of undergraduate instructors who taught introductory life science lab courses. Instructors were invited to participate in this study through three main venues. First, we recruited instructors through email invitations disseminated through CURE-related listservs (e.g., CUREnet) and established CURE programs. Second, we disseminated study information through social media (e.g., Facebook groups, Twitter/X). Finally, instructors were asked to share study information with their colleagues who met the study criteria (i.e., snowball sampling).

Our final analytic sample included 48 instructors who taught introductory biology laboratory courses at 39 unique institutions from 36 states in the United States and Canada (Table 1). These institutions included community colleges, master’s degree-granting universities, and research-intensive universities. Participants included graduate teaching assistants and non-tenured and tenured faculty, and >45% were teaching courses with an enrollment of 20-24 students. On average, the instructors had 7.5 years of teaching experience (SD = 3.0, range = 1-10+). The majority of instructors identified as female (73%) and white (79%). All instructors were provided with a $50 gift card and an individualized report of their results as an incentive for their participation in the study.

**Table 1:** Characteristics of the instructors used in the analyses (*n* = 48).

| **Demographic** | ***n* (%)** |
| --- | --- |
| *Sex* |  |
| Male | 10 (21%) |
| Female | 35 (73%) |
| Other | 3 (6%) |
| *Race/Ethnicity* |  |
| African American or Black | 4 (8%) |
| Asian | 6 (13%) |
| Hispanic or Latinx | 3 (6%) |
| White | 38 (79%) |
| Other | 2 (4%) |
| *Position* |  |
| Graduate student | 8 (17%) |
| Post-doctoral research associate | 1 (2%) |
| Lecturer | 6 (13%) |
| Assistant Professor | 10 (21%) |
| Associate or Full Professor | 12 (25%) |
| Other position* | 11 (23%) |
| *Years of college teaching experience* |  |
| 1-4 years | 7 (15%) |
| 5-9 years | 17 (35%) |
| 10+ years | 24 (50%) |
| *Course size* |  |
| 10-19 students | 8 (16%) |
| 20-24 students | 23 (48%) |
| 25-29 students | 7 (14%) |
| 30+ students | 10 (22%) |
| *Institution type* |  |
| Associate's Colleges | 6 (13%) |
| Baccalaureate Colleges | 3 (6%) |
| Master's Colleges & Universities | 12 (25%) |
| Doctoral Universities | 27 (56%) |

*Note:* \* indicates director, instructional designer, or other instructional positions

#### Students

Undergraduate students who were enrolled in the courses of participating instructors were invited to participate in the study and complete a survey about their experiences in the course. A total of 462 students completed the survey (Table 2). Student respondents were predominantly female (76%), mostly white (51%), in their first year of college (57%), and had conducted research before (63%). Student participants were compensated with a $25 gift card for completing the survey.

**Table 2:**
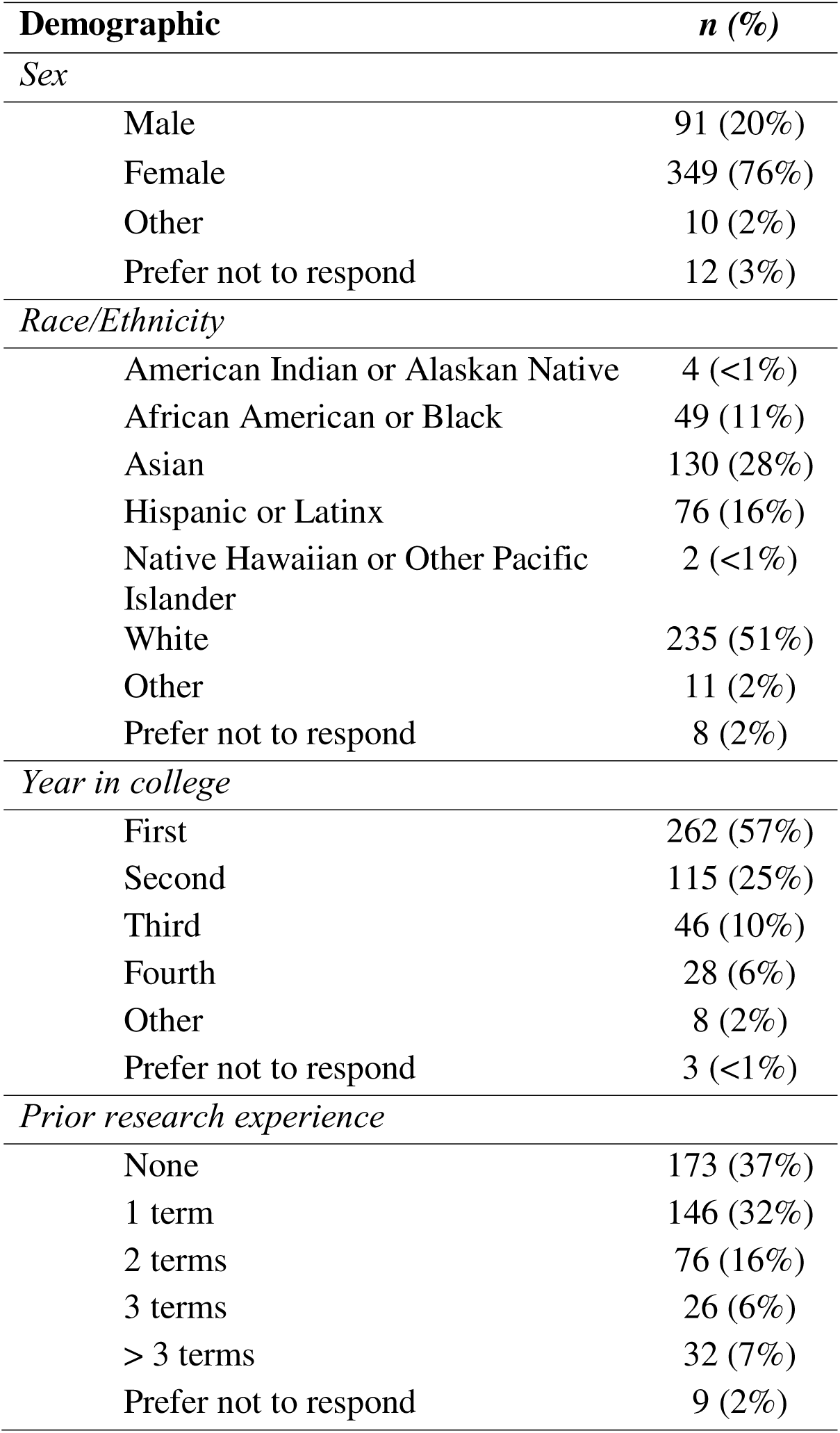
Characteristics of the undergraduates used in the analyses (*n* = 462).

| **Demographic** | ***n* (%)** |
| --- | --- |
| *Sex* |  |
| Male | 91 (20%) |
| Female | 349 (76%) |
| Other | 10 (2%) |
| Prefer not to respond | 12 (3%) |
| *Race/Ethnicity* |  |
| American Indian or Alaskan Native | 4 (<1%) |
| African American or Black | 49 (11%) |
| Asian | 130 (28%) |
| Hispanic or Latinx | 76 (16%) |
| Native Hawaiian or Other Pacific Islander | 2 (<1%) |
| White | 235 (51%) |
| Other | 11 (2%) |
| Prefer not to respond | 8 (2%) |
| *Year in college* |  |
| First | 262 (57%) |
| Second | 115 (25%) |
| Third | 46 (10%) |
| Fourth | 28 (6%) |
| Other | 8 (2%) |
| Prefer not to respond | 3 (<1%) |
| *Prior research experience* |  |
| None | 173 (37%) |
| 1 term | 146 (32%) |
| 2 terms | 76 (16%) |
| 3 terms | 26 (6%) |
| > 3 terms | 32 (7%) |
| Prefer not to respond | 9 (2%) |

### Data Collection

The data for the present study came from two sources: (1) instructor audio recordings of class sessions, which were used to identify instances where instructors used humor, and (2) student surveys about their instructor’s humor and their emotions about the course at the end of the term.

#### Instructor classroom audio recordings

We audio-recorded multiple class sessions during the term for each instructor. Instructors were provided with a lapel microphone and asked to record four class sessions during their course. All instructors recorded their first day of class and three additional class sessions of the instructor’s own choosing. We ultimately collected four class recordings from 90% of instructors (43 out of 48), while 8% (4 instructors) had three class recordings and 2% (1 instructor) had two class recordings. On average, each instructor submitted a total of ~7 hours of classroom recordings (SD = ~2.5 hours, range = 2-12 hours). Recordings were transcribed verbatim and checked for accuracy prior to analysis.

#### Student surveys

We surveyed the students of each instructor using Qualtrics™ at the conclusion of their course. On average, there were 9.5 student responses for each instructor (range = 3-30).

### Measures

A complete list of measures and items are included in the Supplemental Materials.

#### Researcher observations of instructor humor

We used qualitative content analysis^53^ to identify any instances where an instructor used humor. A full description of our content analysis is described elsewhere ^52^, and we describe it briefly here as context for this study with a focus on coding of instructor attempts to be humorous. We began by identifying all possible segments of the text that were considered instructor talk, that is, coherent language used by an instructor that is not directly related to the concepts under study but instead focuses on creating the learning environment ^6^. Multiple coders then independently read each instance of instructor talk identified in all of the transcripts and labelled data from the transcript (e.g., phrases, sentences) that could be considered instances of instructor’s using humor (see Table 3 for examples). We defined instructor use of humor as instances where instructors appeared to make an effort to communicate in ways that could be perceived as funny. After coding the transcripts independently, coders met to discuss any discrepancies to consensus.

**Table 3:** Examples of instructor talk that researchers coded as attempts at humor.

| **Example instance** |
| --- |
| “You can see that my artistic requirements for this course, the bar is very low.” |
| “And if you mess up one cell, it'll destroy your entire bacteria situation going on. And then you'll create a mutant that will destroy the world. No, just kidding.” |
| “I'm a master of awkward silences and once won a staring contest with a goldfish.” |
| “So you guys get the benefit of that, and it's called Shakira. We've named our shaking incubator Shakira 'cuz it shakes.” |

Our analysis was iterative and collaborative. At least four researchers, working in pairs, coded each transcript to consensus. One or more researchers who were undergraduate students at the time of the analyses were involved in coding each transcript for instructor use of humor. The involvement of undergraduate researchers was important for ensuring that we captured all instances of instructor use of humor and for providing a student perspective on the data. The result of our qualitative analysis was a total count of the number of instances of humor for each instructor across all recordings. Given that the length and number of audio recordings varied across instructors, we normalized these values by adjusting them to a per-60-minute basis. Therefore, our final variable for instructor use of humor represents the number of humor attempts made by each instructor per hour.

#### Pleasant and unpleasant emotions

We used a 12-item scale derived from the Circumplex Model of Affect ^54,55^ to measure students’ emotions about their lab courses. Pleasant emotions included elated, enthusiastic, excited, proud, accomplished, and pleased. Unpleasant emotions included stressed, worried, annoyed, bored, apathetic, and exhausted. Students were asked to rate the extent to which each emotion described their affect towards their lab course on a five-point Likert scale (i.e., 1 = “not at all”, 5 = “a great deal”).

We assessed the fit of our four-factor measurement model using confirmatory factor analysis. Our results indicated that the model required modifications, including the removal of two items (apathetic and bored), and the data were best represented using three factors: activated pleasant emotions, deactivated pleasant emotions, and unpleasant emotions (see Supplemental Materials for details). Responses were averaged into a single score for each factor, with higher scores indicating greater levels of emotions. The scale demonstrated acceptable internal reliability (activated pleasant emotions: α = 0.89; deactivated unpleasant emotions: α = 0.88; unpleasant emotions: α = 0.86).

#### Student reports of instructor humor

We asked students to report their perceptions of their instructor’s humor using a single item derived from an instructor-student rapport scale ^56^. Students responded to the item (“My instructor has a good sense of humor”) on a seven-point scale ranging from 1 (strongly disagree) to 7 (strongly agree).

### Analytic Approach

Given the multilevel structure of our data, wherein students are nested within instructors, we tested our hypotheses with linear multilevel modeling. First, we tested the proportion of between instructor variance in each of our student outcomes. The between-instructor variance (i.e., intraclass correlation) ranged from 13-25% for each variable providing evidence of non-negligible variance between instructors. Therefore, we included a random intercept for each instructor in our models.

We next conducted inferential tests to determine the suitably of including random effects at the student level. Specifically, we tested for potential significance of fixed effects in our models by conducting *t*-tests with the Satterthwaite degrees of freedom in order to control for Type I error rates ^57^. Likelihood ratio tests comparing a fixed slopes model to a random slopes model for perceived humor suggested that adding random slopes did not significantly improve the model fit for activated pleasant emotions, χ*^2^*(2) = 5.47, *p* = .065, deactivated pleasant emotions, χ*^2^*(2) = 4.90, *p* = .086, or unpleasant emotions, χ*^2^*(2) = 1.85, *p* = .397. Thus, we retained a random intercept model with a fixed slope for our final analyses.

We fit our linear multilevel models using restricted maximum likelihood (REML) estimation with students (Level 1) nested within instructors (Level 2) using the *lme4* package ^58^ in *R* v4.1.0 in Rstudio (version 2022.12.0+353) ^59^. Following recommendations for centering predictors in multilevel models ^60,61^, we person-mean centered our Level 1 predictor (i.e., student reports of instructor humor) and grand-mean centered our Level 2 predictor (i.e., researcher observations of instructor humor). Our models aimed to understand the effects of both student reports and researcher observations of instructor humor on students’ activated pleasant, deactivated pleasant, and unpleasant emotions about their lab courses. Our models are stated as follows:

Level 1 (within instructors):

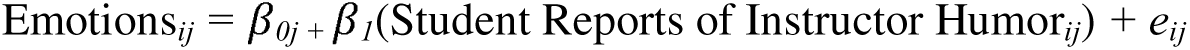

Level 2 (between instructors):

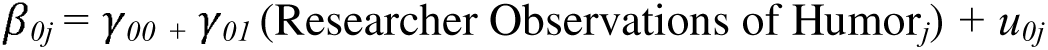

In our Level 1 model, *β_1_* represents the fixed effect of student reports of instructor humor, *β_0j_* is the random intercept, and *e_ij_* represents the residual error for student *i* within instructor *j*’s class. Emotions*_ij_* represents our dependent variable (i.e., student-reported emotions).

In our Level 2 model, *γ_00_* represents the grand mean intercept of student emotions across instructors, *γ_01_* represents the fixed effect of researcher observations of instructor humor (centered at the grand mean), and *u_0j_* represents the random effect for instructor *j*. *β_0j_* represents the average student-reported emotions for instructor *j* after accounting for student-level predictors.

## RESULTS

Means, standard deviations, and correlations of all variables are reported in Table 4. The mean value of researcher observations of instructor humor was 1.39 instances per hour (SD = 1.29 instances) with a range of 0 to 6 instances per hour (Figure 1A). This result suggests that instructor attempts at humor was fairly rare but ranged widely across instructors. In contrast, the mean value of student reports of instructors being humorous was 5.94 (SD = 1.17) on a 7-point scale. This result indicates that, on average, students thought their instructors were quite humorous. For some instructors (e.g., instructors 8, 15, 16, and 19, among others), students’ reports were similar, suggesting high agreement that their instructor was humorous (Figure 1B). For other instructors (e.g., instructors 2, 3, 23, and 48, among others), student reports ranged widely from strongly disagreeing that their instructor was humorous to strongly agreeing (Figure 1B). Consistent with the differing patterns between researcher observations and student reports, we found no correlation between researcher observations and student reports of instructor humor (Table 4).

**Figure 1:**
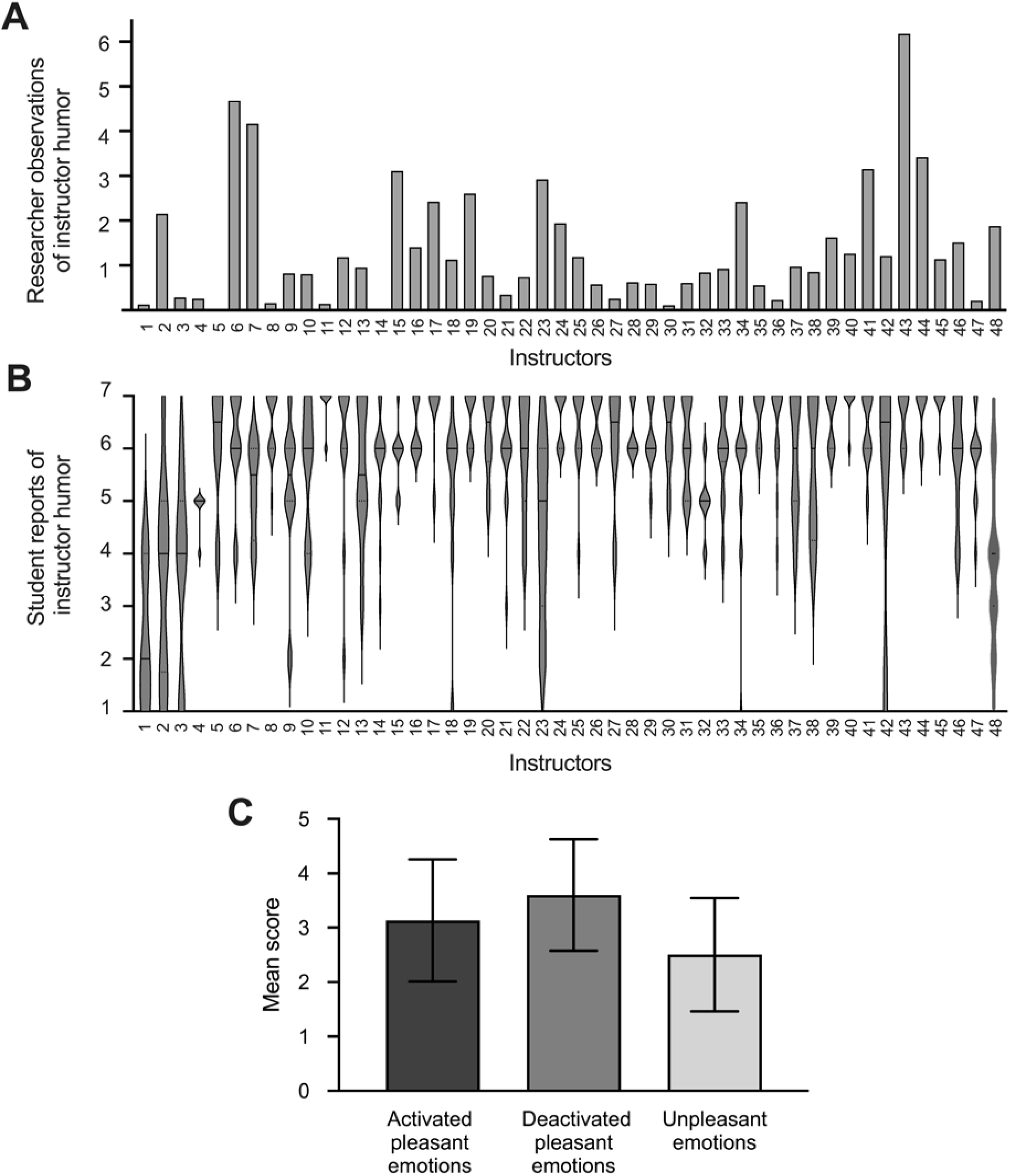
(A) Researcher observations of instructor humor. Units denote average number of researcher-observed instances of instructor humor in a 60-minute period. (B) Student reports of instructor humor for the same instructors as shown in Panel A (1 = strongly disagree, 7 = strongly agree). (C) Overall means and standard deviations for students’ emotions about their lab course (*n* = 462).

**Table 4:** Means, Standard Deviations, & Correlations of Variables. ** *p* < 0.01.

| Variable | *M* | *SD* | *$\alpha$* | Correlations |  |  |  |
| --- | --- | --- | --- | --- | --- | --- | --- |
|  |  |  |  | 1 | 2 | 3 | 4 |
| 1. Researcher observations of instructor humor | 1.39 | 1.17 | - | - |  |  |  |
| 2. Student reports of instructor humor | 5.94 | 1.29 | - | 0.08 | - |  |  |
| 3. Activated pleasant emotions | 3.13 | 1.12 | 0.89 | 0.03 | 0.41** | - |  |
| 4. Deactivated pleasant emotions | 3.60 | 1.02 | 0.88 | 0.03 | 0.40** | 0.78** | - |
| 5. Unpleasant emotions | 2.50 | 1.04 | 0.86 | 0.00 | -0.35** | -0.20** | -0.28** |

On average, students reported pleasant emotions about their lab courses (Figure 1C). Students indicated experiencing both activated pleasant emotions (i.e., elated, enthusiastic, excited; *M* = 3.13, SD = 1.12) and deactivated pleasant emotions (i.e., proud, accomplished, pleased; *M* = 3.60, SD = 1.02) about their courses. Students also reported comparatively lower levels of unpleasant emotions (i.e., stressed, worried, annoyed, exhausted; *M* = 2.50, SD = 1.04).

To address our research questions, we first examined the associations between researcher observations of instructor humor and student emotions. We found that there was no significant association between researcher observations of instructor humor and students’ activated pleasant emotions (*β* = 0.01, *p* = 0.84), deactivated pleasant emotions (*β* = 0.00, *p* = 0.90), or unpleasant emotions (*β* = 0.07, *p* = 0.33) toward their lab courses (Table 5 and Figure 2). These results demonstrate that researcher observations of instructor use of humor are not predictive of students’ emotions about their lab courses.

**Figure 2:**
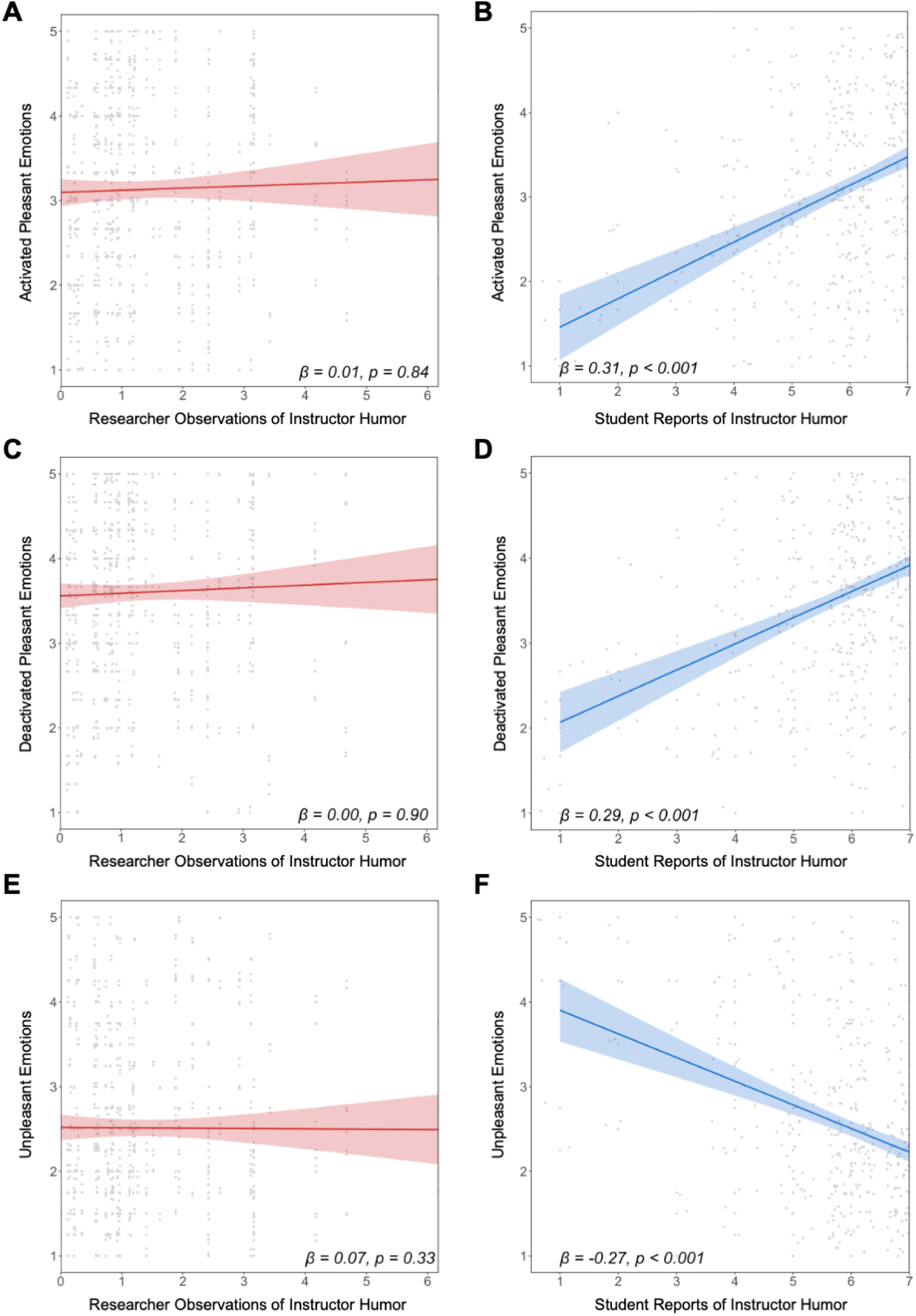
Relationships between researcher observations (red) and student reports (blue) of instructor humor and students’ emotions. Lines represent slopes from multilevel models. Shaded areas represent 95% confidence intervals.

**Table 5:**
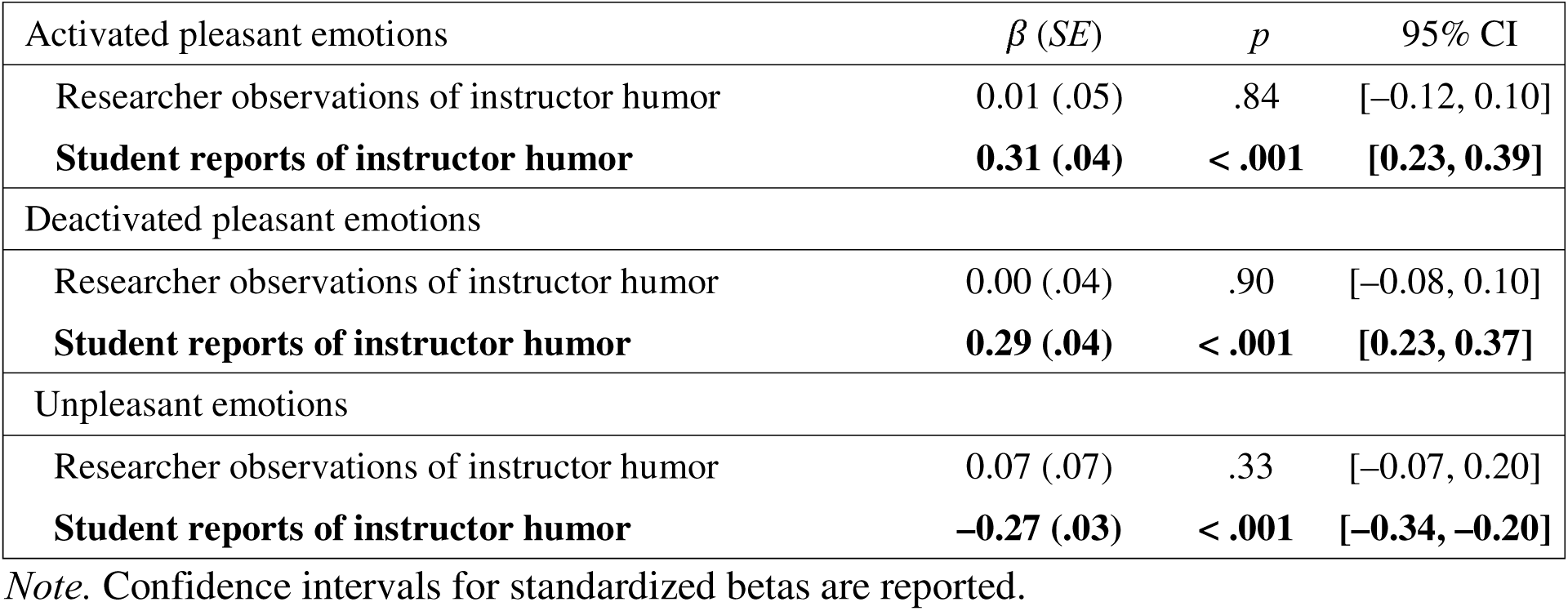
Fixed effects estimates from multilevel models predicting student emotions toward lab course.

In contrast, we found that student reports of instructor humor were significantly associated with their emotions about their lab courses (Table 5 and Figure 2). Students who reported their instructor as being more humorous also reported more activated pleasant emotions (*β* = 0.31 *p* < 0.001), more deactivated pleasant emotions (*β* = 0.29, *p* < 0.001), and fewer unpleasant emotions (*β* = −0.27, *p* < 0.001) toward their lab courses. These findings suggest that students’ perceptions of their instructor’s humor have a medium-sized effect on their emotions about their lab course, while researcher observations of instructor humor do not ^62^.

## DISCUSSION

Students in our study reported more pleasant emotions when their instructor was humorous, including both activated pleasant emotions (e.g., excitement) and deactivated pleasant emotions (e.g., accomplishment). Students also reported more unpleasant (e.g., exhausted) emotions about their lab course when they thought their instructor was less humorous. This result hints at the potential for instructor humor to have positive downstream effects for students, given that students’ emotions predict both their academic behaviors (e.g., study strategies, engagement in class) and their academic performance ^30,33,63^. Future research should test this possibility further by examining the extent to which instructor humor predicts other desirable student outcomes (e.g., learning, academic performance, persistence).

We found that researcher observations of instructor humor were entirely unrelated to student reports of their instructor’s humor. One possible explanation is that we used different data sources for researcher observations and student perceptions. Although researchers analyzed transcripts from a substantial portion of each course (i.e., 25-33% of the 12-13 week courses ^52^) students experienced the full course (100% of class sessions, assuming full attendance). While students rated their instructor’s humor based on the entire term, researchers sampled a limited number of class sessions and the sampling frequency and timing needed to capture instructor’s teaching practices varies ^64^. Furthermore, although people typically forget they are being recorded ^65^ and we chose our recording devices to be as unobtrusive as possible, instructors selected which sessions to record and may have opted to be more serious and less humorous during the class sessions they recorded. Such behavior could reduce the likelihood of researchers observing instructor attempts at humor and undermine our ability to detect a relationship between researcher observations and student reports. In addition, researchers only analyzed the content of the transcripts (i.e., the words instructors used), not the course materials or instructor’s intonations, inflections, timing, or facial expressions. These factors are important for sensing humor, and students had access to these cues. Given the time required for coding instructor humor and the strong predictive power of student reports, researchers should consider using student reports to measure instructor humor in future studies.

Our results also raise interesting questions about measurement of students’ emotions about their lab courses. Although the activated pleasant and deactivated pleasant subscales operated as expected, the unpleasant subscales did not. Specifically, students did not seem to distinguish between activated and deactivated unpleasant emotions, and they appeared to respond differently to the terms “apathetic” and “bored” compared to the other unpleasant emotions. It is possible that “apathetic” is not interpreted as an unpleasant emotion in the same way as the other unpleasant emotion items. Instead, apathy may represent a distinct emotional dimension that is neutral, rather than distinctly pleasant or unpleasant. It is also possible that students weren’t familiar with the term “apathetic,” and it may need to be replaced with a more widely understood term such as “indifferent” or “disinterested.” Our factor analytic results for “bored” indicated that it was its own factor with a modest relationship to “apathetic” and “annoyed.” It may be that students in our sample experienced boredom differently – perhaps as an opportunity to relax cognitively – and thus did not perceive this as an unpleasant emotion. Alternatively, it may be that students in our sample experienced the emotions of “annoyed” and “exhausted” differently than theorized (activated and deactivated, respectively) ^47^.

Future studies should build on the limitations of the present analysis. First, we measured student perceptions of instructor humor and emotions at a single timepoint, which limits our ability to draw conclusions about causality. It may be that students begin to experience more pleasant emotions and less unpleasant emotions as their course progresses and then they view their instructor in a more favorable light (i.e., emotions leading to humor perceptions), instead of humor perceptions predicting emotions. Future research should survey students about instructor humor and emotions across multiple time points and use more sophisticated analytic methods to better understand their relationship over time. Second, our results relied on a single survey item to measure instructor humor, which may limit measurement quality. Future studies should examine whether multi-item scales, such as the Humor Style Questionnaire ^66^, provide insights into the influence of different humor types (e.g., self-defeating, affiliative or relationship building) and offer greater predictive utility.

Our results also have implications for teaching. Mainly, instructors should consider using humor during lab instruction. Although being funny may seem simple, using it in a course may require expert knowledge on how and when to use it effectively ^67^. Instructors may benefit from learning about uses of humor. Instructors could reflect on published descriptions of appropriate and inappropriate uses of humor to consider what they feel comfortable trying ^68^. Instructors may also benefit from reading reviews of effective uses of humor in classrooms, including recommendations from experts in humorous communication ^41^, and discussing with colleagues how they could apply this knowledge in their teaching. Importantly, students in our study who were in the same course did not always agree that their instructor was humorous. Thus, more in-depth investigation into the range of ways students perceive humor in lab courses (e.g., through more sophisticated measurement) and instructor reflection on what may or may not work for them and their students in their setting are important next steps for maximizing the potential benefits of humor.

## Acknowledgements

We thank our participants for sharing their time and perspectives. We also thank the undergraduate members of the Vertically Integrated Projects Program for their contributions to the analyses and the Social Psychology of Research Experiences & Education lab for their feedback on our results. This material is based upon work supported by the National Science Foundation under award number 2021138. Any opinions, findings, and conclusions or recommendations expressed in this material are those of those author(s) and do not necessarily reflect the views of the National Science Foundation.

